# Blind Virtual Screening at Scale: A Scalable End-to-End Pipeline for Blind Docking and Affinity Prediction

**DOI:** 10.1101/2025.10.10.681617

**Authors:** Matthew Parks, Xin Yu, Yuxing Peng, Jayant Bhambhani, Darren Hsu, Duminda Ranasinghe, Emine Kucukbenli, Hok Hei Tam

## Abstract

Accurate and scalable prediction of protein–ligand interactions remains a central challenge in computational drug discovery, especially when the binding site is unknown (i.e., blind docking). We present a high-throughput, end-to-end algorithm for virtual screening that combines DiffDock, a diffusion-based generative model for blind docking, with UniDock Vina, an algorithm for rapid scoring. We benchmarked this approach on the CASF-2016 and DUD-E datasets, analyzing pose quality, scoring accuracy, and screening performance. We find that competitive screening power can be achieved when generating and scoring as few as three poses and without pose refinement, which facilitates scalability. Notably, our method achieves 86.78% and 82.00% for the percent of actives among the top 1% and 10% of ranked ligands, respectively, when generating as few as three poses per protein-ligand pair. The workflow is scalable, supporting blind docking and affinity prediction at a mean throughput of 0.76 seconds per protein-ligand pair when generating 40 ligand poses in batched mode parallelized to 8 NVIDIA A100 80G GPUs. These results demonstrate that accurate, large-scale blind virtual screening is feasible and offers a practical solution for screening against novel or less characterized protein targets. Code is available at: https://github.com/xinyu-dev/blind-screening-benchmark

## 1. INTRODUCTION

Estimation of the potential for a binding interaction between a protein and a ligand is a critical component of many drug discovery campaigns. At fine resolution, insights into the mechanism of action of a candidate drug, as well as its medicinal properties, can be obtained from examination of the contacts between a docked ligand and the protein. At a coarser resolution, virtual screening of candidate compounds for a target protein can be facilitated by high-throughput computational predictions of protein–ligand interactions.

There are several complex steps required for estimation of a potential protein-ligand interaction, including identification of a potential binding pocket on the target protein, docking of the candidate ligand to the binding pocket to obtain the ligand pose, and estimation of the binding affinity (scoring). In the local docking scenario, the binding pocket of the target protein and perhaps even the ligand pose are known. However, in large-scale virtual screening scenarios— particularly early-stage or exploratory ones—neither the binding pocket nor pose are available. Blind docking may be an option for these cases, where ligands are docked to the entire protein surface without a predefined binding pocket. The quality of interaction estimation depends on both the accuracy of the docked ligand pose and the robustness of the scoring algorithm.

Recent advances in machine learning (ML) models for blind docking, such as DiffDock (*1*) and TankBind (*2*), have demonstrated substantial improvements in pose prediction accuracy, generalization to unseen protein targets, and computational efficiency. These models can outperform traditional physics-based models in throughput and flexibility, making them promising tools for structure-based drug discovery without predefined binding pockets. In particular, DiffDock—a diffusion-based generative model—has demonstrated strong performance for blind docking to fixed protein structures (*1*).

However, the potential for binding interaction must be estimated for a ligand pose. Binding affinity, representing the interaction strength between a protein and ligand for a putative ligand pose, could be used to estimate the potential for binding interaction. Algorithms that approximate physics-based scoring rules can facilitate higher-throughput estimation of binding affinity from specified ligand poses.

Recent efforts have sought to adapt these models for end-to-end virtual screening. For instance, Fortela et al. (*3*) scored poses generated by DiffDock with smina, using smina binding affinity scores (*4*) and DiffDock confidence scores in a focused case study of four perfluoroalkyl and polyfluoroalkyl substances (PFAS). Wu et al. (*5*) introduced a deep learning model that simultaneously folds, docks, and scores binding affinity, but its high computational burden inhibits scalability, and the study did not report screening power. Similarly, Morehead et al. (*6*) developed a benchmarking pipeline for blind docking algorithms that evaluates scoring power across datasets but does not assess screening power, a key metric for virtual screening.

To address this gap, we developed a scalable, end-to-end algorithm for blind docking and affinity prediction that combines generative pose modeling via DiffDock with efficient physics-based scoring via UniDock Vina. Our implementation leverages NVIDIA’s BioNeMo (*7*) to provide optimized, containerized environments for deploying and scaling models like DiffDock on GPU infrastructure. The following sections describe the design of our algorithm, the experimental setup for benchmarking, and the results of our evaluation across standard benchmark datasets.

## 2. METHODS

To evaluate the performance, scalability, and practical applicability of our pipeline for blind virtual screening, we conducted a series of experiments using two standard benchmark datasets: CASF-2016 and DUD-E. These experiments were designed to assess the contributions of each component: pose generation, scoring, and batching. We also sought to determine the overall performance of our method in terms of screening power, scoring accuracy, and computational efficiency.

### 2.1 Benchmarking with the CASF-2016 Dataset

The CASF-2016 dataset (*8*) is a curated collection of 285 protein:ligand pairs with quantitative experimental binding affinities as log K_a_. In this dataset, all protein-ligand pairs were true binders. We benchmarked three independent trials of 20 configurations of our algorithm on the CASF-2016 dataset of 285 protein-ligand complexes, spanning 17,100 computational docking experiments.

For each of the 285 protein-ligand complexes in the CASF-2016 dataset, we generated *P* poses from DiffDock. We repeated this experiment three times for each of *P* = {5, 10, 20, 40}. Thus, the total number of pose samples was 64,125 = 285 x (5 + 10 + 20 + 40) x 3.

All experiments were conducted using a single NVIDIA A100 80G GPU in the NVIDIA DGX cloud environment. The DiffDock component used the DiffDock (legacy version: bionemo_diffdock_nim:24.03.04) container from BioNeMo NVIDIA Inference Microservice (NIM) while scoring was performed with UniDock Vina (version 1.1). For the CASF-2016 experiments, batched docking mode (many ligands to one protein) was not used, and each protein-ligand complex was processed separately.

### 2.2 Benchmarking with the DUD-E Dataset

For the virtual screening benchmarks, we used the standard DUD-E (*Database of Useful Decoys – Enhanced*) dataset (*9*), which includes 102 protein targets. Each protein target is associated with two types of ligands:

● Actives: molecules known to bind the protein
● Decoys: molecules with similar properties to actives (e.g., molecular weight, logP), but not expected to bind

SMILES strings (*10*) were extracted for actives and decoys. For each SMILES, if at least one conformer could be generated by RDKit (*11*), it was considered as a valid SMILES.

For each target, active ligands were combined into a single SDF file, and decoys were combined into a separate single SDF file.

Protein-ligand complexes were sub-selected among all possible complexes in DUD-E as follows:

1. Include all 102 protein targets from DUD-E.
2. Let N_j_ be the number of active ligands for protein target *j*.
3. Then select N_j_ /2 active ligands for each protein target *j* for inclusion in our benchmark.
4. For each protein target *j*, randomly select N_j_/2 decoys from the large set of decoys for protein target *j* for inclusion in our benchmark.
5. The final dataset used for our benchmark contained 102 targets and a total of 22,754 ligands.

DiffDock was invoked on the DUD-E dataset via the NVIDIA BioNeMo (*7*) NIM (version 1.2, same weights as bionemo_diffdock_nim:24.03.04, but with the ‘many ligands to one protein’ batched docking mode). We used batched docking (many ligands per protein) and SMILES input.

Calculation of the enrichment factor (EF) was performed through bootstrapping, with a starting decoy:active ratio set at 50 for each target (58.09 due to rounding). A total of 2000 independent bootstrap sampling trials were conducted.

### 2.3 UniDock Vina

To score binding affinity for DiffDock-generated poses, we used UniDock Vina with a dynamic docking box that included a buffer of ±5 Å in all three dimensions (x, y, z). Because each ligand pose may occupy a different spatial position, we used paired batch mode, which allowed individual specification of the docking box location and dimensions for each ligand.

Although paired batch mode is slower than GPU batch mode, it was necessary when ligands in a batch required distinct box configurations. In contrast, GPU batch mode supported only a single, uniform box per batch of ligands.

We evaluated four UniDock Vina scoring configurations across different experiments:

#### No refinement ("Local Only" Mode)

● Energy minimization using an automatic docking box (±4 Å in each dimension)
● CPU-only execution
● Multiprocessing, used to parallelize scoring tasks

#### Detailed Mode

● Used exhaustiveness = 512 and max_step = 40
● Balances computational efficiency with scoring accuracy

#### Refinement Mode

● Used exhaustiveness = 1024 and max_step = 40 for more detailed pose refinement
● Required preparing a docking box configuration for each pose
● Box size was defined as ligand dimensions with a ±5 Å margin

These different modes allowed us to explore trade-offs between runtime performance and scoring precision across our virtual screening experiments.

### 2.4 Evaluation Metrics

To evaluate the performance of our algorithm, we used three complementary metrics: scoring power, ranking power, and screening power, with each capturing a different aspect of predictive effectiveness.

● Scoring Power was measured as the Pearson correlation coefficient between the predicted binding affinities generated by our algorithm and the experimentally determined binding affinities (reported as log K_a_ values) from the CASF-2016 dataset. This metric reflected the model’s ability to reproduce the actual magnitude of binding affinity. This metric was computed across targets.
● Ranking Power was computed using the Spearman rank correlation coefficient between predicted and experimental affinities, analogously to scoring power. This measure evaluated how well the model preserved the order of affinities across compounds. This metric was computed per target and then summarized across targets by averaging.
● Screening Power assessed the ability of our method to distinguish active ligands from decoys. Using the DUD-E dataset, we benchmarked performance by calculating enrichment factors (EFs) for the top α = 1% and 10% of ranked compounds for each target. We conducted bootstrap sampling (n = 2000 trials) with a fixed decoy-to-active ratio of 58:1. The enrichment factor at threshold α was computed as:

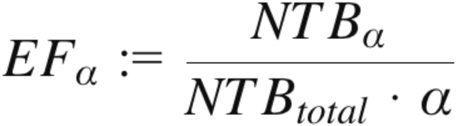

where *NTB_α_* is the number of true binders among the top α-percent (e.g. α = 1% or 10%) of ranked compounds, and *NTB_total_* is the total number of true binders for a given target protein.

## 3. RESULTS

We hypothesized that a scalable, performant method for blind virtual screening could be derived from the synthesis of a model for blind docking and a model for scoring binding affinity. In particular, we developed a method that estimated the binding affinity between a ligand and a protein by generating ligand poses for a fixed protein structure without specification of a binding pocket (i.e., blind docking), scoring the binding affinity of predicted ligand pose(s), and finally summarizing the scores across poses. The blind docking step was based on DiffDock; the scoring step was based on UniDock Vina.

We optimized and evaluated the performance of our virtual screening algorithm using two benchmark datasets: CASF-2016 (*8*) and DUD-E (*9*). We characterized performance via screening power, scalability, ranking power, and scoring power, where screening power and scalability were the primary metrics for our use case of blind virtual screening.

### 3.1 Method Design

One approach for predicting the binding affinity between a ligand and a protein is to predict the binding affinity of candidate ligand poses obtained from docking the ligand to the protein.

Docking a ligand to a protein without prior specification of a binding pocket or ligand pose is termed blind docking. Towards this end, we constructed a method based on a pre-trained DiffDock model for blind docking and on UniDock Vina for scoring binding affinity (**Figure 1**).

**Figure 1:**
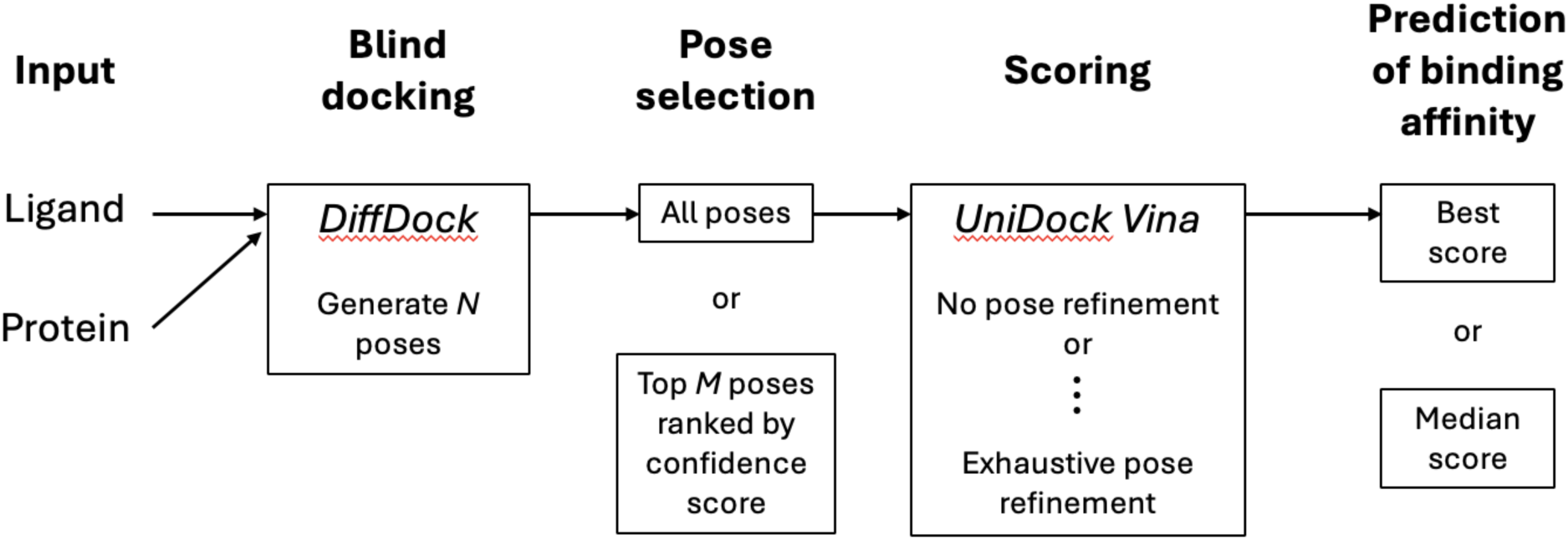
General design of our algorithm and variations explored. *The input was a SMILES for a ligand and a PDB for a protein. Poses were generated from DiffDock. Poses from DiffDock were then scored by UniDock Vina. An estimate of binding affinity was derived from the scores for the poses*.

DiffDock is a generative ML model that stochastically generates ligand poses for a given fixed protein structure. We used the trained DiffDock model published by Corso 2023 (*1*), accelerated by NVIDIA BioNeMo (*7*). Each ligand was represented by a SMILES string (*10*). The 3D structure of the target protein was provided as a Protein Data Bank (PDB) file.

A critical parameter was the number of ligand poses to generate for a single protein-ligand pair. Generating more poses may increase the probability of finding a high-quality pose, but at the cost of lower computational throughput. With the CASF-2016 dataset, we generated 5, 10, 20, or 40 poses per ligand from DiffDock. With the DUD-E dataset, we generated 40 poses per ligand from DiffDock. Given a ligand pose from DiffDock, we used UniDock Vina to score the putative binding affinity. UniDock Vina can optionally refine the ligand pose. Refinement can range from minimal refinement with the “local only” mode to substantial pose remodeling by running Unidock’s full pocket-specific docking command. With the CASF-2016 dataset, we extracted the top pose based on DiffDock confidence score, and applied Unidock’s “local only” mode to minimally refine, and score the pose with Unidock’s built-in vina function. For the DUD-E dataset, we compared Unidock’s “local only” mode against its docking refinement mode with exhaustiveness set to 1024 and max_step set to 40, followed by scoring by the same Vina function.

Another important step was the derivation of a single assessment of interaction potential from the multiple poses and associated scores. For the CASF-2016 dataset, we selected the pose that had the highest DiffDock confidence score as the representative pose. For the DUD-E dataset, we explored several approaches:

1. Retain the top five valid DiffDock poses ranked by confidence score, then score each with Unidock, and select the pose with the best Vina score.
2. Retain all valid DiffDock poses (up to 40), score each with Unidock, and select the pose with the best Vina score.
3. From all valid poses, perform 2000 bootstrap sampling trials with N = 1, 3, 5, 10, 20, or 30 poses. Score each with Unidock and either use the median Vina score or select the pose with the best Vina score.

A summary of the above experiments is shown in Figure 1.

### 3.2 A Pan-Protein Threshold for DiffDock Confidence Scores Is Likely Infeasible

DiffDock produces a confidence score for each stochastically generated pose. The DiffDock confidence score predicts the probability that the root mean square distance (RMSD) of the generated pose is less than 2 Å. A procedure for selecting poses generated by DiffDock for subsequent scoring could involve filtering poses by their assigned confidence scores. The utility of such an approach would depend on the sampling variance of confidence scores and their distribution across proteins.

To investigate the behavior of DiffDock confidence scores as a function of number of generated poses and target protein, we generated DiffDock poses for all 285 protein-ligand pairs in the CASF-2016 dataset and examined the distribution of confidence scores. To assess the potential of obtaining a positive confidence score for a pose in a protein-ligand complex, we counted how many complexes had at least one pose with a positive confidence score (i.e., >50% probability of RMSD < 2 Å), repeating this process for three independent trials. We found that the number of protein-ligand complexes with a positive confidence score for the top-ranked pose increased with the number of poses generated per protein-ligand complex (**Figure S1**), suggesting that a pose with a positive confidence score could be attained with sufficiently large sampling of poses per protein-ligand complex. Further, the best observed DiffDock confidence score among sampled poses was largely consistent across independent trials and across different number of poses sampled per trial (**Figure S2)**, suggesting that a procedure based on filtering by DiffDock confidence scores would not suffer from high sampling variance even when sampling as few as five poses, thereby supporting the scalability of our method.

A threshold for the DiffDock confidence score of poses that is general, rather than protein-specific, could support virtual screening for proteins with few or no well-characterized binders, or for high-throughput screening across many proteins. To investigate the potential for selecting a general, pan-protein threshold for the DiffDock confidence score, we compared the distributions of confidence scores across the 57 proteins in the CASF-2016 dataset. We found that the distribution of confidence scores varied systematically by protein (**Figure S3)**, thereby frustrating the prospect of selecting a single, pan-protein threshold for DiffDock confidence scores for filtering generated poses.

### 3.3 DiffDock Confidence Scores are Not a Sufficient Proxy for Binding Affinity

Despite the systematic variation in DiffDock confidence scores across proteins, we hypothesized that if DiffDock confidence scores were strongly correlated with experimental properties, especially binding affinity, then filtering poses by the DiffDock confidence score before scoring with UniDock Vina could improve overall algorithm performance. A threshold for DiffDock confidence scores specific to each protein or complex could be used if DiffDock confidence scores correlate with complex resolution. To the contrary, we found a weak positive correlation between DiffDock confidence score and experimental complex resolution (**Figure S4**, **Figure S5**). Alternatively, if DiffDock confidence scores correlated with binding affinity, then further scalability of our algorithm could be achieved by filtering poses for each protein or complex by confidence score prior to subsequent scoring with UniDock Vina. We observed positive but weak correlation between DiffDock confidence scores and experimental binding affinities in the CASF-2016 dataset (**Figure S6**). In particular, the pose with the best DiffDock confidence score often did not have the best UniDock Vina score (**Figure S7**).

### 3.4 Scoring Power Increases with Number of Poses

A simple and least computationally burdensome configuration would be to score only the single pose with the best confidence score from DiffDock. To estimate the performance of this approach, we sampled 5, 10, 20, or 40 poses from DiffDock for each of the 285 protein-ligand pairs from CASF-2016, scored the pose with the best confidence score with UniDock Vina, and then computed the scoring power and ranking power against the experimental binding affinities. We found that scoring power increased with the number of poses from 0.38 for 5 poses to 0.48 (∼28% increase) for 40 poses (**Table 1**). In contrast, ranking power did not change significantly (**Table 1**), suggesting that the top-ranked DiffDock pose was robust across the number of poses in terms of ranking power, but it was necessary to sample more poses to improve scoring power.

**Table 1:** Scoring power and ranking power versus number of generated poses. For each protein-ligand complex in CASF-2016, 5, 10, 20, or 40 poses were generated by DiffDock, and the pose with the best DiffDock confidence score was subsequently scored with UniDock Vina. Mean and standard deviation values were derived from three independent trials. Runtimes were recorded on a single NVIDIA A100 80G GPU.

| Number of poses generated | Mean scoring power (± standard deviation) | Mean ranking power (± standard deviation) | Mean runtime per protein-ligand pair (seconds) |
| --- | --- | --- | --- |
| 5 | 0.376 ± 0.049 | 0.396 ± 0.036 | 3.4 |
| 10 | 0.366 ± 0.09 | 0.398 ± 0.016 | 3.8 |
| 20 | 0.439 ± 0.051 | 0.42 ± 0.013 | 5.3 |
| 40 | 0.48 ± 0.016 | 0.402 ± 0.024 | 7.9 |

### 3.5 Scalability by Batching Ligands per Protein Target

In a high-throughput blind virtual screening scenario, many compounds are docked to the same protein. Towards scalable virtual screening, we found that the accelerated version of DiffDock from NVIDIA BioNeMo scales with the number of poses, with mean runtime per pose decreasing from 0.68 seconds per pose when generating 5 poses to 0.20 seconds per pose when generating 40 poses (**Table 1**).

Further, we explored several computational strategies for batching many ligands for docking against the same protein, available through the BioNeMo implementation of DiffDock. We found that batching protein-ligand complexes into batches of 16 ligands per protein obtained the best throughput (**Table 2**).

**Table 2:** Runtime for different batching strategies. Times represent the protein in PDB “7rwo” from DUD-E dataset.

| Method | Trial 1 (total seconds) | Trial 2 (total seconds) | Trial 3 (total seconds) | Mean runtime (total seconds) | Mean time (seconds) per protein-ligand pair |
| --- | --- | --- | --- | --- | --- |
| Batch all 32 ligands and submit as 1 query | 42.1 | 43.6 | 43.7 | 43.1 | 1.35 |
| Split 32 ligands into 2 batches (16 ligands/batch), run sequentially | 43.5 | 41.6 | 44.6 | 43.2 | 1.35 |
| Split the 32 ligands into 2 batches (16 ligands/batch), using asyncio | 41.3 | 41.1 | 42.7 | 41.7 | **1.30** |
| Split the 32 ligands into 2 batches (16 ligands/batch), and using multiprocessing | 41.5 | 41.7 | 42.9 | 42.0 | **1.31** |

### 3.6 Scalability and Protein Sequence Length

We docked 22,754 ligands against 102 proteins from the DUD-E dataset with DiffDock. For each of the 102 protein targets, we partitioned the ligands into 8 batches and generated 40 poses per protein-ligand pair. Each batch was parallelized to an A100 GPU, in a cluster with 8 A100 GPUs. The mean runtime per pose was 0.76 seconds. We found that average prediction time per ligand correlated with protein length, and using larger batch sizes reduced the computational time per ligand (**Figure 2**).

**Figure 2:**
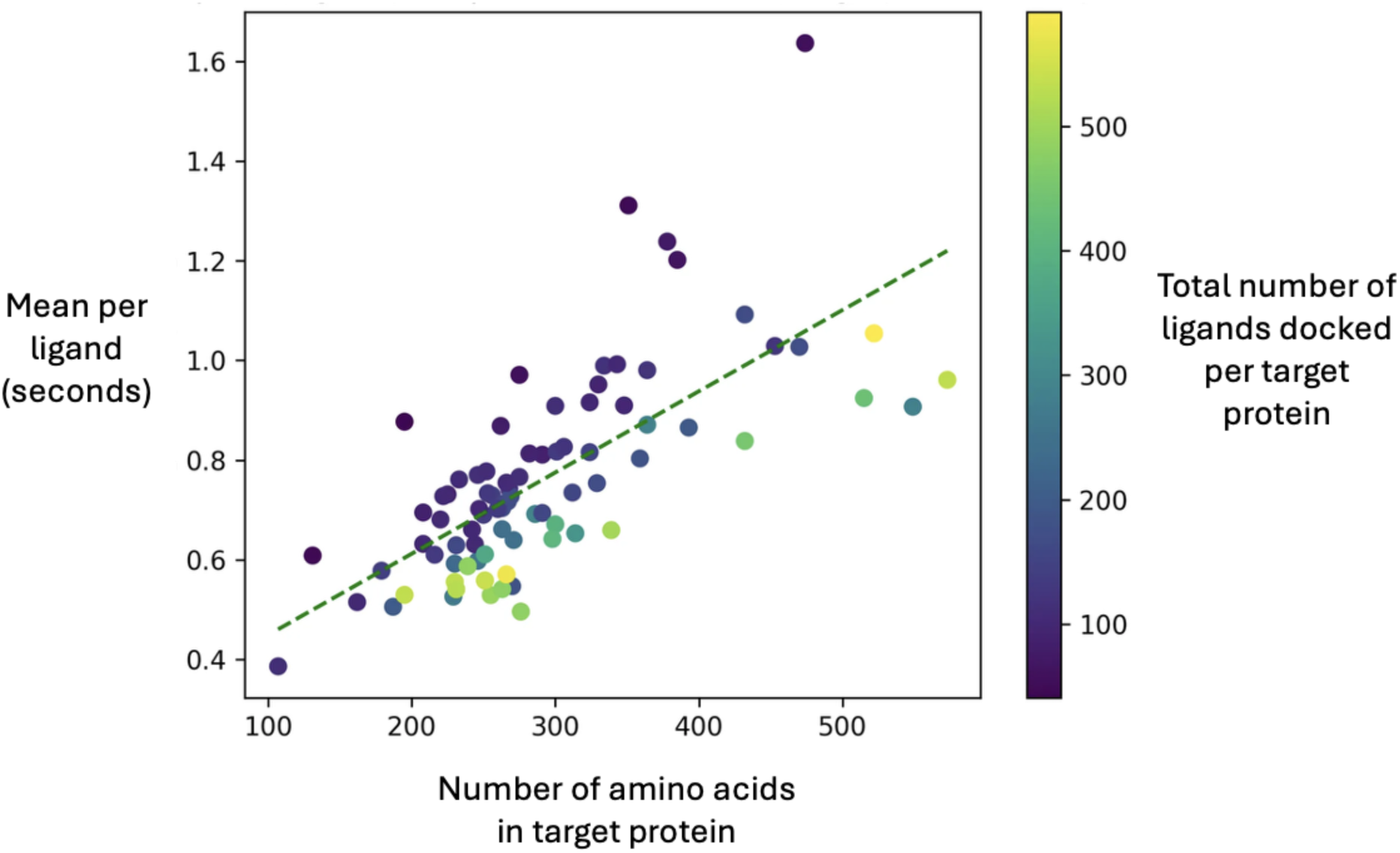
Runtime per protein-ligand pair. *The x-axis is the length of the protein in a protein-ligand complex. The y-axis represents the mean compute time (in seconds) per ligand. Each point represents one of the 102 protein targets in the DUD-E dataset. Points are colored by the total number of ligands (active and decoys) for each target. For every target, we divide the ligands into 8 batches of equal size, therefore the color of the dots (total number of ligands) also correlates with batch size*.

### 3.7 Comparison of Screening Power to Local Docking Models

Screening power represents the capacity of an algorithm to distinguish binders from non-binders (also termed actives vs. inactives) and is a crucial metric for a high-throughput virtual screening algorithm. The DUD-E dataset, spanning 102 protein targets, provides a challenging benchmark for screening power as decoy compounds are computationally generated for each protein target to have similar chemical properties to the known actives.

We evaluated several variants of our method by using different methods for selecting poses, scoring poses with UniDock Vina, and summarizing scores. Namely, we explored combinations from among the following variants: 1) generating different numbers of poses from DiffDock; 2) selecting the top 5 poses ranked by DiffDock confidence score for subsequent scoring with UniDock Vina; 3) scoring all generated poses and using the best Vina score; 4) scoring all poses and using the median of the Vina scores.

For each algorithm variant under consideration, we estimated the binding affinity for each protein-ligand complex in DUD-E with our algorithm and ranked the ligands per protein target by predicted binding affinity. We evaluated screening power through several metrics: 1) the percent of protein targets in which the ligand with the best Vina score is an active; 2) the percent of actives in the top 1% of ligands per protein (“Top 1%”); 3) the percent of actives in the top 10% of ligands per protein (“Top 10%”); 4) enrichment factor for the top 1% of ligands; and 5) enrichment factor for the top 10% of ligands.

Surprisingly, scoring poses with UniDock Vina without any pose refinement outperformed exhaustive pose refinement across all metrics for screening power (**Table 4**). This result further supports the scalability of the algorithm, as scoring without refinement is substantially faster than with refinement.

**Table 4:** Screening power for variants of our method based on the top-5 poses ranked by DiffDock confidence score. . *For each protein-ligand complex from DUD-E, 40 poses were generated with DiffDock. The top 5 poses ranked by DiffDock confidence score were then scored with UniDock Vina either without refinement (“local only” mode, executed on CPUs) or with refinement (executed on GPUs). The best UniDock Vina score across the 5 selected poses per protein-ligand complex was declared to be the predicted binding affinity score for the protein-ligand complex. We evaluated screening power performance by ranking ligands per protein target by predicted binding affinity and calculating the percentage of actives in the top 1% and top 10% of ranked compounds. “Bootstrap EF” refers to enrichment factors estimated from 2,000 bootstrap trials. Brackets indicate 95% confidence intervals across the trials. Timings were for a compute node with 8x A100 GPUs and 240 CPU cores*.

| Pose refinement mode | Percent of Actives in single best-scored ligand | Percent of actives in top 1% of ligands per protein | Bootstrap EF for in top 1% of ligands per protein | Percent of actives in top 10% of ligands per protein | Bootstrap EF for in top 10% of ligands per protein | Mean runtime (seconds per ligand) |
| --- | --- | --- | --- | --- | --- | --- |
| No refinement | 83% | **83.56**<br>[77.08, 90.05] | **11.83**<br>[11.72, 11.94] | **79.59**<br>[75.72 - 83.47] | **3.29**<br>[3.28, 3.31] | **<0.01** |
| Refinement | 68% | 73.66<br>[65.90 - 81.42] | 8.23<br>[8.13, 8.32] | 73.33<br>[68.83 - 77.83] | 2.76<br>[2.75, 2.78] | 0.3 |

When filtering by the top five poses ranked by DiffDock confidence score, we found that the percent of active ligands in the top 1% and top 10% of ligands ranked by predicted binding affinity per protein target was 83.56% and 79.59%, respectively, with enrichment factors of 11.83 and 3.29, respectively (**Table 4**). For comparison, a baseline model would yield 50% actives in the top 1%, given the 1:1 active-to-decoy ratio per target in the initial dataset (See Methods).

### 3.8 Competitive Screening Power is Sustained for Limited Poses

Our method achieved an enrichment factor of 15.05 (95% bootstrap CI [14.93, 15.17]) for the top 1% and 3.67 (95% bootstrap CI [3.66, 3.69]) for the top 10% of ligands when generating and scoring 40 poses. The screening power achieved by our method based on blind docking is competitive with, or outperforms, other algorithms for which the binding pocket was predefined (*12*) on the DUD-E dataset (**Table 5**), though our estimates are derived from a random subset of the protein-ligand pairs in the DUD-E dataset due to computational limitations However, the competitive screening power we found for 40 poses per protein-ligand complex comes at the cost of scalability. We investigated the effect of reducing the number of poses per protein-ligand complex on screening power. Towards this end, we bootstrap resampled 1, 3, 5, 10, 20, or 30 poses from the 40 generated poses. Performance of our method dropped only slightly when generating as few as 3 poses, achieving top 1% and top 10% active rates of 86.78 and 82.00, respectively (**Table 6**). Further, we found that the percent of active ligands in the top 1% of ranked ligands was still 86.34% when sampling even just a single pose from DiffDock, compared to 87.99% when sampling 40 poses (**Table 6).** This result further enhances the scalability of the algorithm, as the computational burden is substantially less when generating 3 poses as opposed to 40 (**Table 1**).

**Table 5:** Comparison of enrichment factors to published algorithms. *Enrichment factors for UniDock and AutoDock are derived from Yu, et al, 2023* (*12*).

| Algorithm | Docking scenario | Pose refinement mode | Enrichment Factor for Top 1% of ligands per protein |
| --- | --- | --- | --- |
| UniDock | local | fast | 11.1 |
| UniDock | local | balanced | 12.0 |
| UniDock | local | detailed | 13.1 |
| AutoDock-GPU | local | fast | 2.6 |
| AutoDock-GPU | local | balanced | 2.8 |
| AutoDock-GPU | local | detailed | 3.1 |
| Vina-GPU | local | fast | 11.9 |
| Vina-GPU | local | balanced | 13.5 |
| Vina-GPU | local | detailed | 12.3 |
| Ours | blind | no refinement | 15.0 |

**Table 6:** Screening power for variations of our method based on scoring all generated poses. . *For each protein-ligand complex from DUD-E, 40 poses were generated with DiffDock. All poses were scored by UniDock Vina without refinement. Generating N = 1, 3, 5, 10, 20, 30 poses was simulated by bootstrap resampling from amongst the set of 40 generated poses per protein-ligand complex. The best UniDock Vina score across resampled poses per protein-ligand complex was declared to be the predicted binding affinity score for the protein-ligand complex. We evaluated screening power performance by ranking ligands per protein target by predicted binding affinity and calculating the percentage of actives in the top 1% and top 10% of ranked compounds. Brackets indicate 95% confidence intervals across all trials*.

| Method to select poses generated by DiffDock | Percent of actives among the best ligand for each protein | Percent of actives in Top 1% of ligands per protein | Percent of actives in Top 10% of ligands per protein |
| --- | --- | --- | --- |
| Use all valid poses ($\leq 40$ per each PL complex) | **89%** | **87.99**<br>[82.21, 93.77] | 82.24<br>[78.44, 86.04] |
| Bootstrap sample N=1 poses | 86.42<br>[86.29, 86.54] | 86.34<br>[86.23, 86.45] | 79.91<br>[79.87, 79.95] |
| Bootstrap sample N=3 poses | 86.55<br>[86.43, 86.66] | 86.78<br>[86.68, 86.88] | 82.00<br>[81.97, 82.04] |
| Bootstrap sample N=5 poses | 86.60<br>[86.49, 86.71] | 86.71<br>[86.62, 86.81] | 82.24<br>[82.21, 82.27] |
| Bootstrap sample N=10 poses | 86.93<br>[86.83, 87.03] | 86.58<br>[86.49, 86.66] | **82.38**<br>**[82.35, 82.41]** |
| Bootstrap sample N=20 poses | 87.33<br>[87.25, 87.41] | 86.29<br>[86.22, 86.37] | 82.22<br>[82.19, 82.24] |
| Bootstrap sample N=30 poses | 87.83<br>[87.76, 87.91] | 86.38<br>[86.31, 86.44] | 82.21<br>[82.19, 82.23] |

### 3.9 Using the Median-scored Pose Improves Screening Power

In the above experiments, we used the best binding affinity scored by Uni-Dock Vina amongst all considered poses from DiffDock. We hypothesized that this approach is less robust to outliers or edge-cases in the generation or scoring of poses. Alternatively, we compared performance when using the median score from Uni-Dock Vina amongst the considered poses from DiffDock.

We found that the percent of active ligands in the top 1% and top 10% of compounds ranked by binding affinity score from Uni-Dock Vina significantly and substantially improved when using the median score across poses instead of the best scores among poses (**Table 7**). In particular, using the median score across just three poses improved top 1% actives to 88.27%, compared to 86.78% when using the single best score (**Table 7)**.

**Table 7:** Enrichment factors for screening power using the lowest versus median score from UniDock Vina among poses. *Results are shown for the top 1% and 10% of ligands per target*.

| Score selection strategy | Number of poses per bootstrap resample | Percent of Active Ligands Among Top 1% of Ligands | Percent of Active Ligands Among Top 10% of Ligands |
| --- | --- | --- | --- |
| Lowest | 3 | 86.78 [86.68, 86.88] | 82.00 [81.97, 82.04] |
| Median |  | 88.27 [88.17, 88.37] | 81.57 [81.53, 81.60] |
| Lowest | 5 | 86.71 [86.62, 86.81] | 82.24 [82.21, 82.27] |
| Median |  | 89.29 [89.19, 89.38] | 82.20 [82.17, 82.23] |
| Lowest | 10 | 86.58 [86.49, 86.66] | 82.38 [82.35, 82.41] |
| Median |  | 90.13 [90.03, 90.22] | 82.79 [82.76, 82.82] |
| Lowest | 20 | 86.29 [86.22, 86.37] | 82.22 [82.19, 82.24] |
| Median |  | 90.76 [90.67, 90.84] | 83.25 [83.22, 83.28] |
| Lowest | 30 | 86.38 [86.31, 86.44] | 82.21 [82.19, 82.23] |
| Median |  | 91.08 [91.00, 91.16] | 83.43 [83.41, 83.45] |

## 4. DISCUSSION

High-throughput blind virtual screening can substantially accelerate the drug discovery process and increase the likelihood of success. We have described a scalable end-to-end method for blind virtual screening based on blind docking, not requiring specification of a binding pocket or a ligand pose, that performs competitively with or outperforms state-of-the-art algorithms that *do* require specification of the binding site in terms of screening power.

Our method achieved a trade-off between scalability and screening power on the one hand and scoring power and ranking power on the other, which is favorable for the virtual screening application. Specifically, we have shown that our method achieved competitive or state-of-the-art screening power, is applicable to the blind docking setting, and is simultaneously highly scalable. Subsequent efforts could endeavor to improve ranking power and/or scoring power without significantly sacrificing scalability or screening power, which are both useful for virtual screening.

Biases in the construction of datasets used to benchmark machine learning models can compromise the reliability of estimates of model generalization. Studies have highlighted such biases for the DUD-E dataset and demonstrated discrepancies in generalization as a result (*13,14*). Although our method was not trained on these datasets, such biases frustrate the comparison of performance estimates across benchmark datasets.

A key limitation of our approach is that it requires a protein structure. Crystal structures of only a fraction of the proteins in the human proteome are available in public databases like RCSB PDB, and the appropriate protein structure may be different per biological context. One approach could be to generate candidate structures for proteins using models like AlphaFold or ESMfold, though performance would need to be characterized for this approach.

Another limitation is the assumption of our method on a fixed protein structure. Recent advances in simultaneous prediction of protein structure with an interacting bound ligand, such as with Boltz-1 (*15*), present the opportunity for relieving these assumptions and improving all metrics—screening power, ranking power, and scoring power— though at the cost of reduced scalability. Batched processing, for example with BioNeMo from NVIDIA, for Boltz-1 may reduce the impact on scalability in this trade-off.

Our results facilitate virtual screening for hundreds of thousands of candidate compounds against many proteins at computationally reasonable costs and time. This capacity facilitates screening of compounds for on-target activity, but also for filtering out compounds that have undesirable off-target interactions. Owing to the scalability and efficiency of batch processing, our algorithm could be used for filtering out compounds based on off-target effects against a large set of off-target proteins relevant to a particular pathway.

## 5. CONCLUSION

We present an end-to-end algorithm for high-throughput virtual screening that combines blind docking via a generative model for pose prediction (DiffDock) with affinity prediction via a fast, physics-inspired scoring method (Uni-Dock Vina). Without relying on prior knowledge of the binding site or ligand pose, our method achieved competitive screening power across a large benchmark (DUD-E), with a Top 1% actives rate of 86.78%.

Crucially, our method scales efficiently to tens of thousands of protein–ligand pairs on a single GPU, and performance remains high even when scoring only a few generated poses. Median-based scoring selection further enhances robustness, mitigating the impact of pose selection bias. These features make the approach well suited for early-stage virtual screening, particularly when structural information is incomplete or unavailable.

By demonstrating that generative docking combined with lightweight scoring can achieve competitive performance with traditional structure-based methods, this work offers a practical and scalable framework for blind virtual screening against novel or understudied targets.

## 6. SUPPLEMENTARY FIGURES

**Figure S1:**
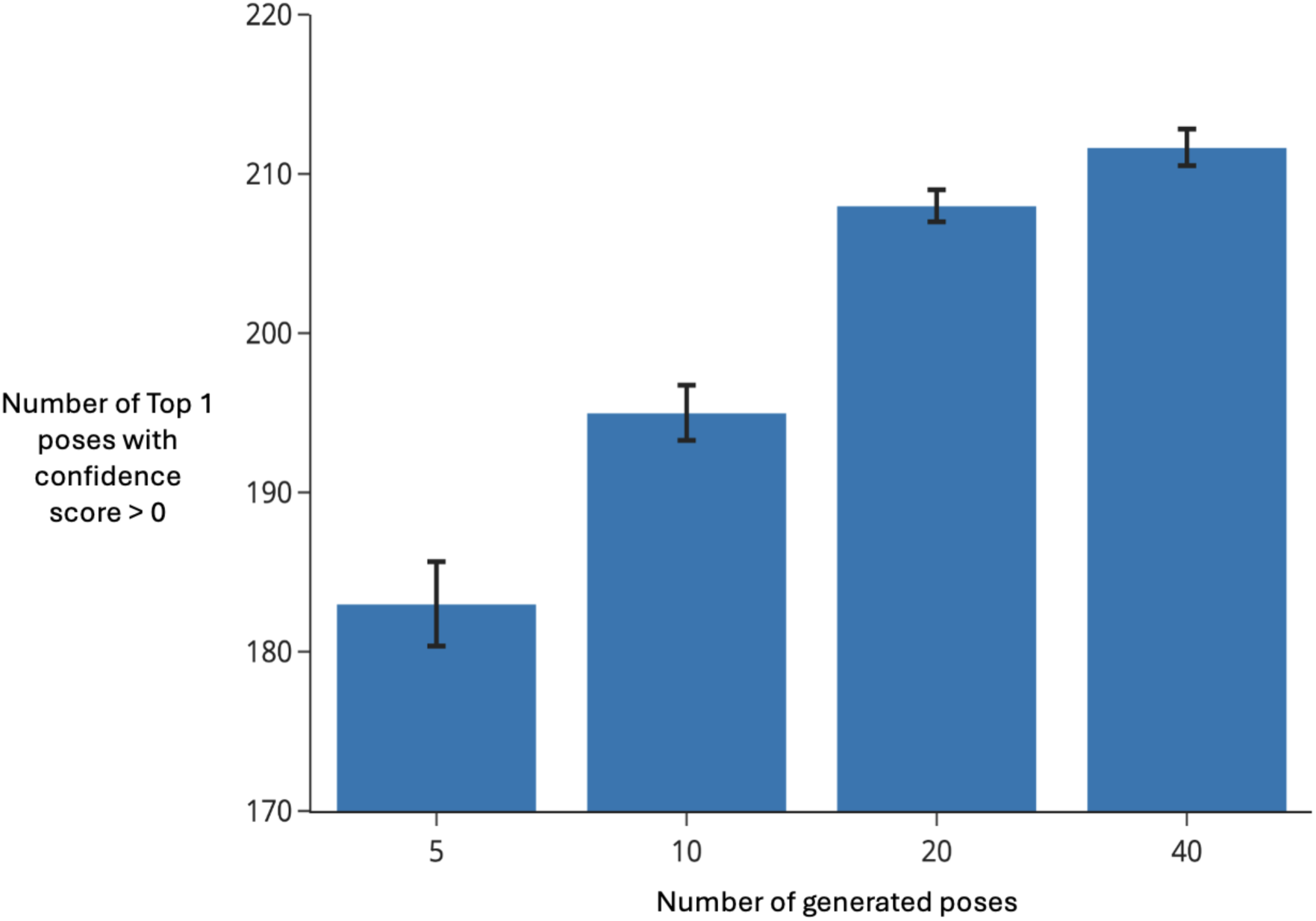
**The best DiffDock confidence score as a function of number of poses generated by DiffDock for CASF-2016**. *Number of protein:ligand complexes with a best DiffDock confidence score > 0, as a function of the number of poses generated. For each of the 285 CASF-2016 complexes, 5, 10, 20, or 40 poses were generated across 3 independent trials. Bars show the mean number of complexes with a top 1 confidence score > 0 ; error bars represent ±1 standard deviation among the three trials*.

**Figure S2:**
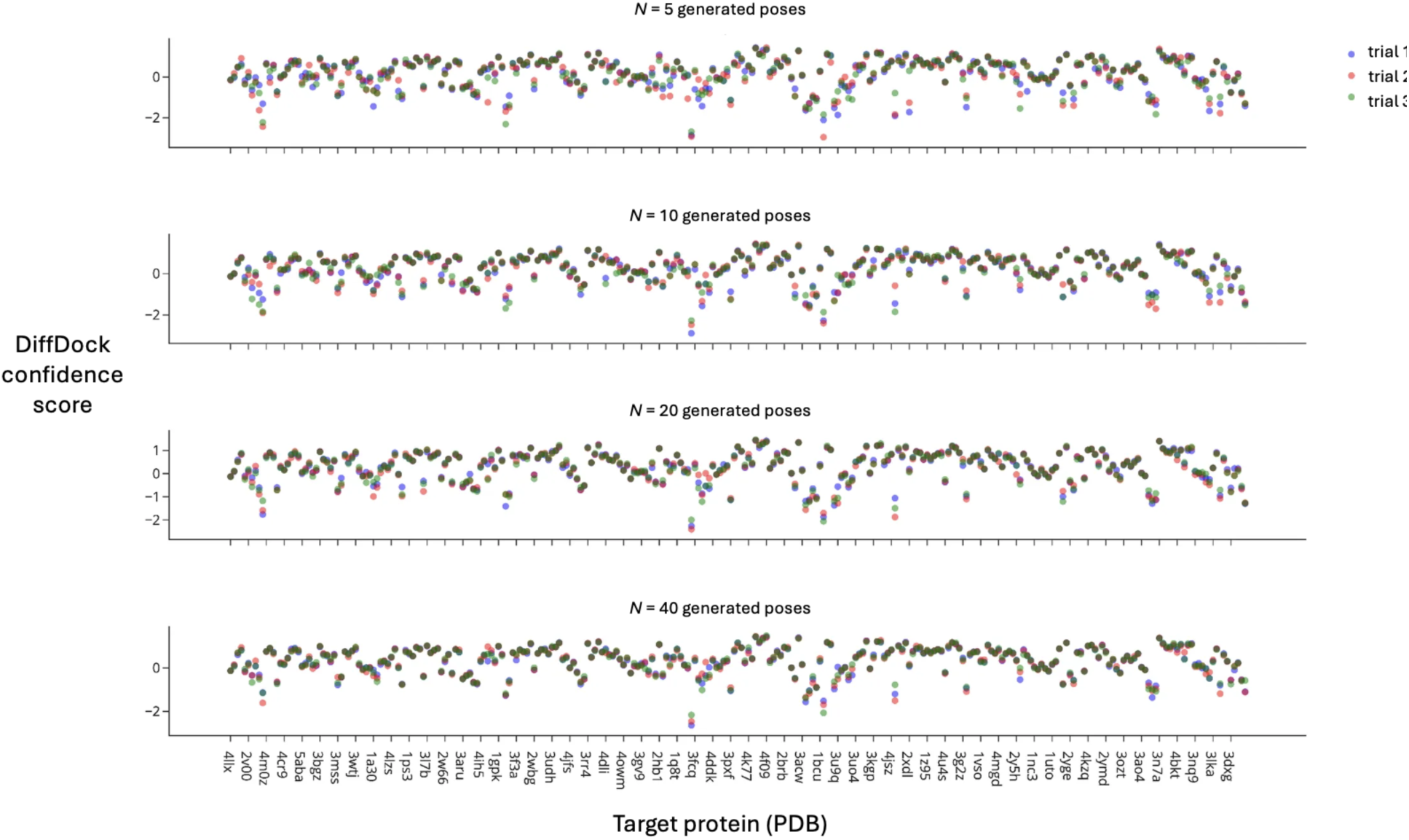
**Variance of DiffDock confidence scores across 3 independent trials**. *Each color represents an independent trial of DiffDock. In each trial, 5, 10, 20, or 40 poses were generated for each of the 285 protein:ligand pairs in CASF-2016. The 57 distinct protein targets in the CASF-2016 dataset are represented along the x-axis. The y-axis gives the best DiffDock confidence score among the generated DiffDock poses per protein:ligand pair*.

**Figure S3:**
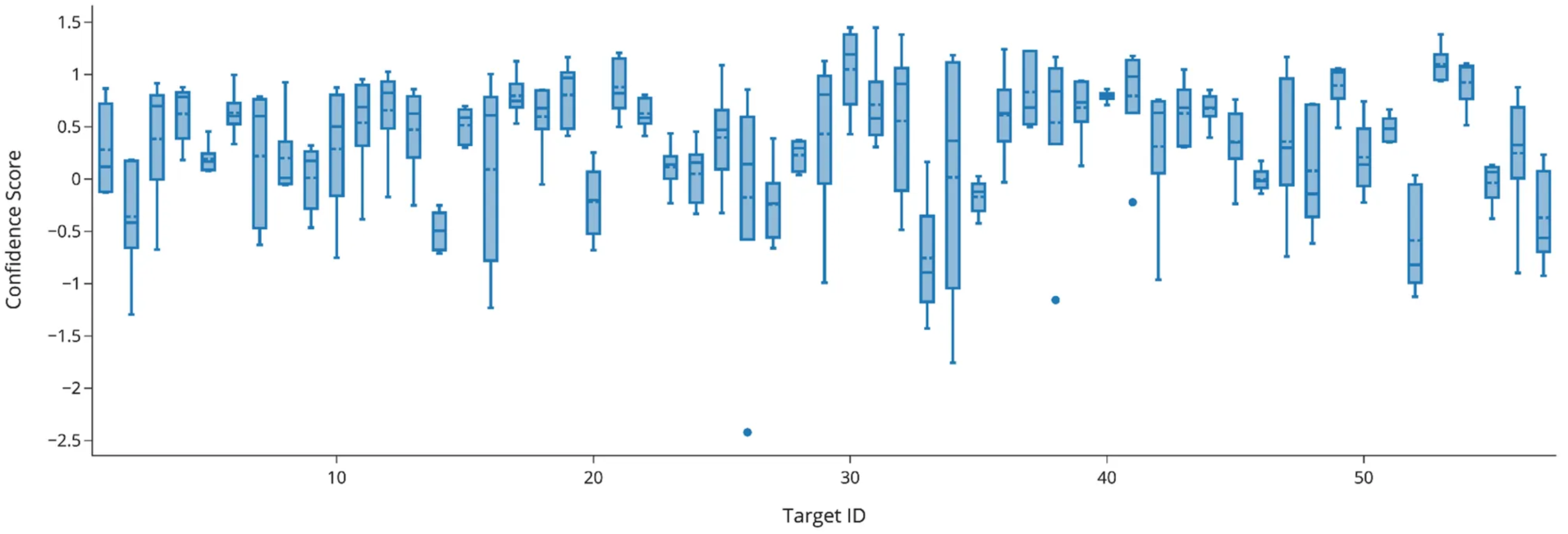
Distribution of DiffDock confidence scores by protein target. Protein targets are arranged along the x-axis. The y-axis represents the mean of the best DiffDock confidence scores across 3 independent trials for a protein-ligand complex. In each trial, 40 poses were generated per protein-ligand complex and the best DiffDock confidence score among the 40 poses per protein-ligand complex was selected. Box plots represent the distribution of these summary statistics across ligands for the same protein target.

**Figure S4:**
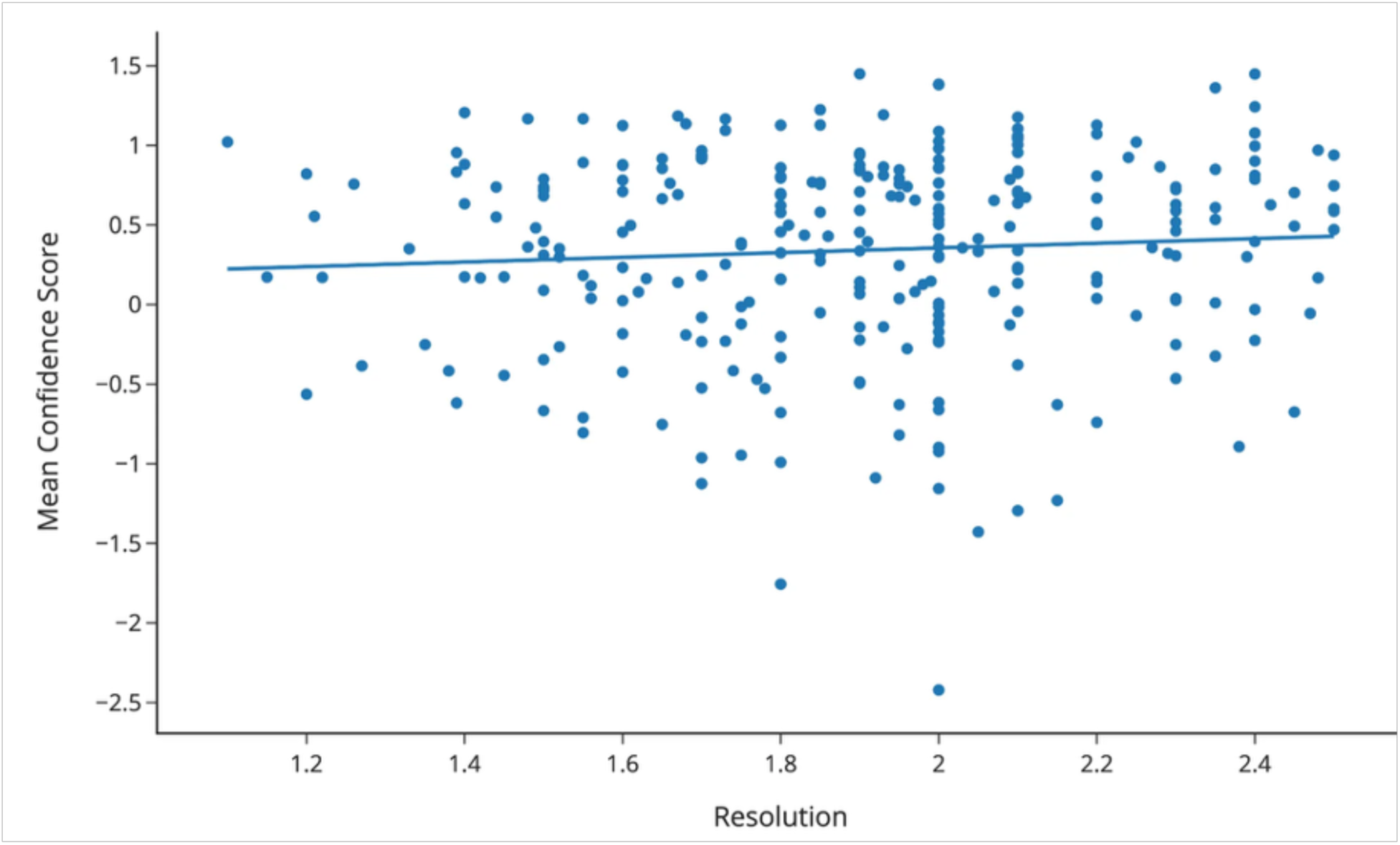
DiffDock confidence scores versus complex resolution. Each point represents one of the 285 protein:ligand complexes in the CASF-2016 dataset. The x-axis gives the true resolution of the protein:ligand complexes in CASF-2016. The y-axis represents the mean across three independent trials of the *best DiffDock confidence scores from 40 generated poses*.

**Figure S5:**
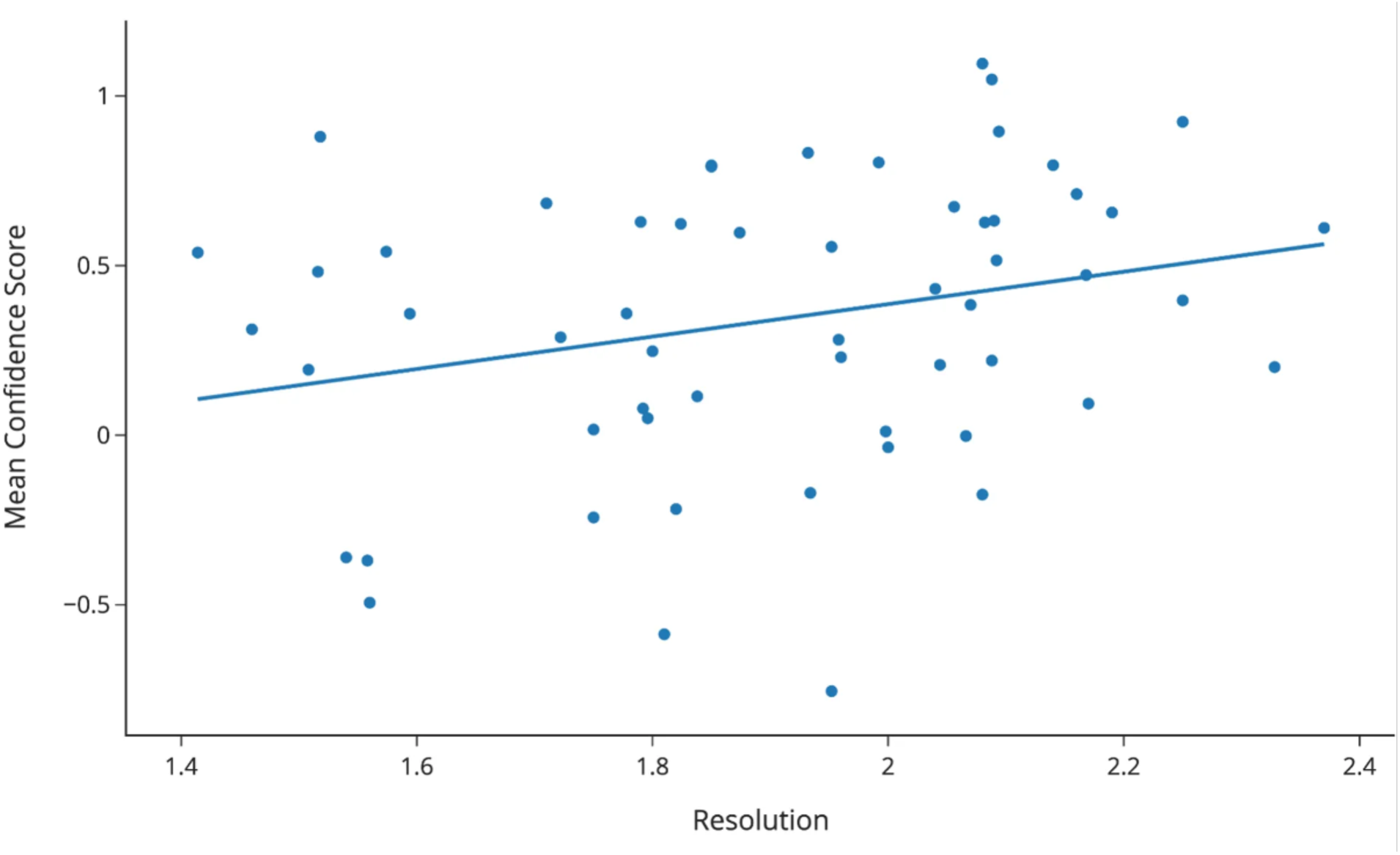
DiffDock confidence scores versus complex resolution per protein target. Each point represents one of the 57 distinct proteins in the CASF-2016 dataset. The x-axis gives the mean resolution (Angstrom) among the five PDB structures per protein. The y-axis gives the mean of the best DiffDock confidence scores across the 3 independent trials of generating 40 poses for each of the 285 protein:ligand complexes. Pearson correlation coefficient r = 0.266.

**Figure S6:**
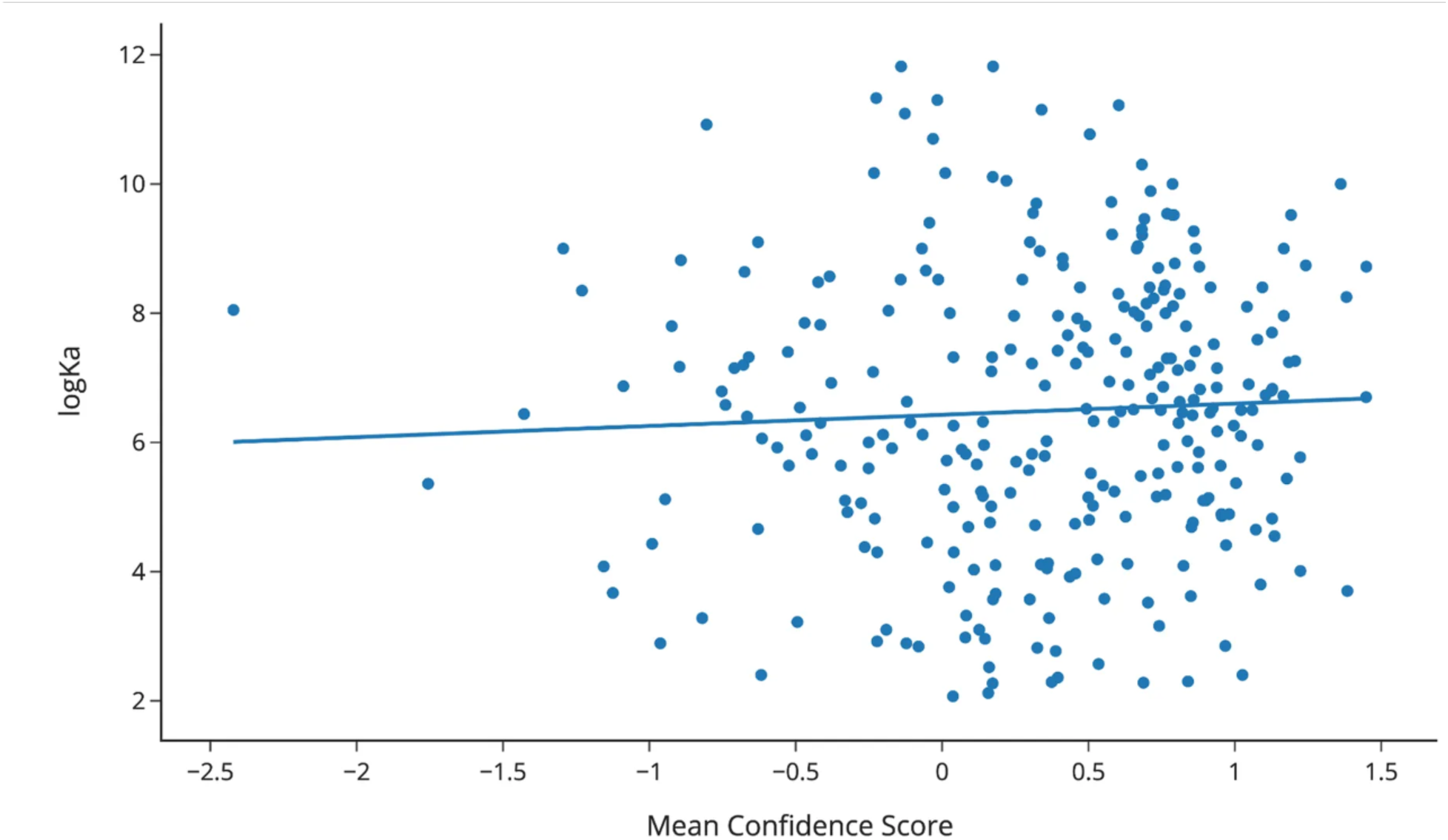
**DiffDock confidence scores versus experimental binding affinity**. *The y-axis represents experimental binding affinity in log Ka units, as provided by CASF-2016* (*1*)*. The x-axis gives the mean of the best observed DiffDock confidence scores across 3 independent trials of generating 40 poses for each of the 285 protein:ligand complexes in CASF-2016. Each point represents one of the 285 protein:ligand complexes in the CASF-2016 dataset. Pearson correlation coefficient r = 0.05*.

**Figure S7:**
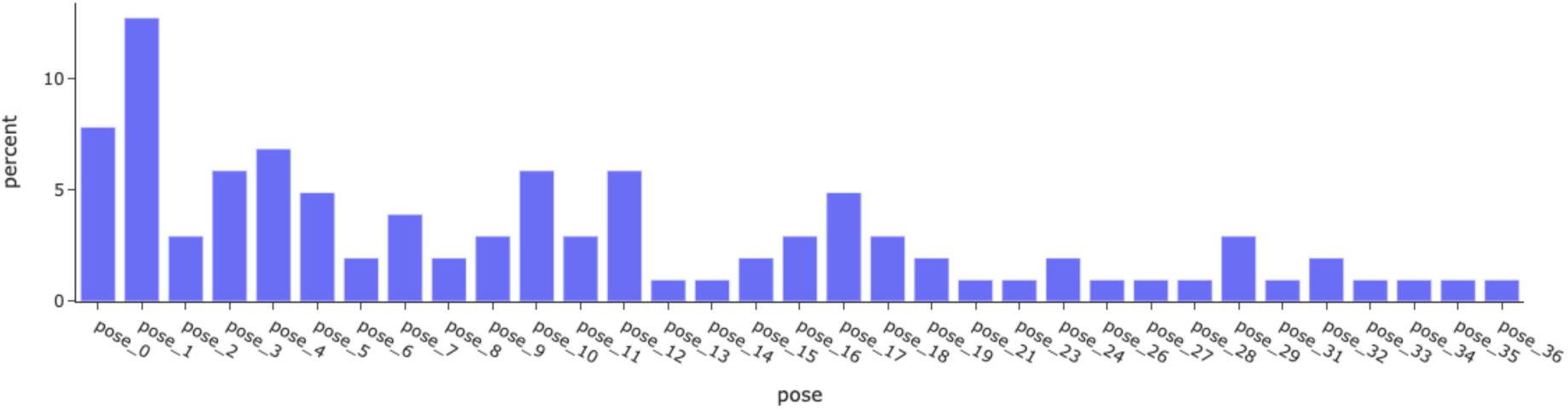
Poses ranked by DiffDock confidence score versus the best binding affinity predicted by UniDock Vina. For each of the 102 protein targets in DUD-E, we selected the compound with the best UniDock Vina score. The x-axis represents the ranked pose from DiffDock across 40 poses for the best compound per the 102 protein targets in DUD-E, ranked by DiffDock confidence score. The pose with the best DiffDock confidence score is ranked as pose 0, and the worst as pose 39. The y-axis gives the fraction of the 102 protein targets in DUD-E for which the ranked pose (by DiffDock confidence score) has the lowest binding affinity score predicted by UniDock Vina among the poses.

